# Interpretable Deep Learning for Breast Cancer Cell Phenotyping Using Diffraction Images from Lens-Free Digital In-Line Holography

**DOI:** 10.1101/2021.05.29.446284

**Authors:** Tzu-Hsi Song, Mengzhi Cao, Jouha Min, Hyungsoon Im, Hakho Lee, Kwonmoo Lee

## Abstract

Lens-free digital in-line holography (LDIH) offers a wide field of view at micrometer-scale resolution, surpassing the capabilities of lens-based microscopes, making it a promising diagnostic tool for high-throughput cellular analysis. However, the complex nature of holograms renders them challenging for human interpretation, necessitating time-consuming computational processing to reconstruct object images. To address this, we present HoloNet, a novel deep learning architecture specifically designed for direct analysis of holographic images from LDIH in cellular phenotyping. HoloNet extracts both global features from diffraction patterns and local features from convolutional layers, achieving superior performance and interpretability compared to other deep learning methods. By leveraging raw holograms of breast cancer cells stained with well-known markers ER/PR and HER2, HoloNet demonstrates its effectiveness in classifying breast cancer cell types and quantifying molecular marker intensities. Furthermore, we introduce the feature-fusion HoloNet model, which extracts diffraction features associated with breast cancer cell types and their marker intensities. This hologram embedding approach allows for the identification of previously unknown subtypes of breast cancer cells, facilitating a comprehensive analysis of cell phenotype heterogeneity, leading to precise breast cancer diagnosis.

## Introduction

Breast cancer presents considerable challenges due to its substantial inter-tumor and intra-tumor heterogeneity [1]. In clinical practice, determining the hormone and growth factor receptor status of breast cancer tissues is crucial for effective treatment, as it significantly influences the progression and phenotypes of breast cancer [1–6]. Specifically, the measurement of nuclear estrogen (ER) or progesterone receptors (PR) levels helps determine the responsiveness of breast cancer cells to anti-estrogen therapies such as tamoxifen, fulvestrant, or aromatase inhibitors. Additionally, the presence of human epidermal growth factor receptor 2 (HER2) on the surface of breast cancer cells is a crucial factor, with HER2-positive cancers benefiting significantly from anti-HER2 treatment involving Herceptin [1–4, 6].

Based on the hormone status, breast cancer can be classified into four types: ER/PR- HER2-, ER/PR-HER2+, ER/PR+HER2+, and ER/PR+HER2-, enabling physicians to determine appropriate treatment strategies [3]. Furthermore, recent advancements in molecular and gene expression studies have led to the identification of various breast cancer subtypes, which have implications for tailored clinical treatments [4]. Achieving precise diagnosis and analysis of these breast cancer types or subtypes from heterogeneous tissue samples can enable more efficient and effective treatment for breast cancer patients. However, the current diagnostic workflow based on light microscopes faces limitations such as limited data throughput and high costs, impeding the accurate diagnosis of breast cancer.

Lens-free digital in-line holography (LDIH) is a powerful imaging technique that encodes the 3D information of an object into a single shot of 2D diffraction patterns, i.e., holograms, and computationally converts them to object images [7–8]. LDIH produces images of wide field-of-view (FOV) at micrometer-scale resolution, which cannot be achieved by conventional lens-based microscopes. Because holograms can be obtained to extract 3D cell morphological and structural information in a wide FOV [6], it provides a much larger amount of high-resolution image data than lens-based microscopes of similar FOV. It was also used to monitor and recognize real-time microorganisms to investigate their dynamic behaviors [7–9]. In addition, LDIH has a deep field of depth for digitally refocusing to different reconstruction planes, which can overcome the technical limitations of extracting 3D morphological characteristics of samples in comparison to lens-based microscopes [10–11]. While LDIH has been used in various cellular analyses [12–14] and object tracking [15–16], the conventional workflow of LDIH includes image reconstruction using phase retrieval algorithms, requiring substantial computational resources [17–20] and incurring image artifacts or information loss.

Deep learning (DL) offers a promising solution by bypassing the costly image reconstruction step and directly processing raw holograms. DL excels at recognizing meaningful and often hidden features within complex datasets, making it highly effective for analyzing diffraction images that may not be readily interpretable by human intuition [21–24]. This obviates the need for reconstruction processing, eliminating potential errors or artifacts associated with the reconstruction step. As a result, DL-based methods offer robust applications for LDIH, significantly enhancing the reliability and accuracy of various analyses across multiple domains. These include but are not limited to image reconstruction [25–26], detection of micro-environmental pollution [27], as well as classification and monitoring of diverse biological samples for disease diagnosis [6, 28–31]. Rivenson *et al*. [26] used a deep learning model to perform holographic reconstruction using multi-height phase retrieval in hologram images. O’Connor *et al.* developed deep learning models for cell and disease identification [30, 31]. Recently, DL has been applied to raw holograms for classification tasks. Min *et al.* developed an artificial intelligence diffraction analysis (AIDA) platform for high-throughput cancer cell analysis using LDIH, where deep neural networks performed the recognition/classification of breast cancer cells from raw holograms [6]. Kim *et al.* have used a transfer learning approach to directly classify raw holograms generated from cells and microbeads without a reconstruction process [32].

Most of these existing applications of deep learning methods primarily focus on standard image translation and classification tasks, leaving the utilization of raw holograms for other machine learning tasks such as regression or unsupervised learning largely unexplored. Moreover, explaining how deep learning methods analyze raw holograms is crucial as direct application of human intuition is not feasible in this context. Unfortunately, the current landscape lacks a deep learning model that can deliver accurate interpretability for raw holograms. In this paper, we introduce HoloNet, a novel deep learning architecture designed to offer superior performance in both supervised and unsupervised learning tasks, while also providing enhanced interpretability for the analysis of LDIH holograms, as illustrated in **Figure 1**. Our proposed model enabled precise classification of diverse breast cancer cell types and accurate estimation of marker intensity values from raw hologram. Furthermore, through the utilization of feature-fusion HoloNet, we successfully identify previously unknown cell phenotypes within the context of breast cancer.

**Figure 1.**
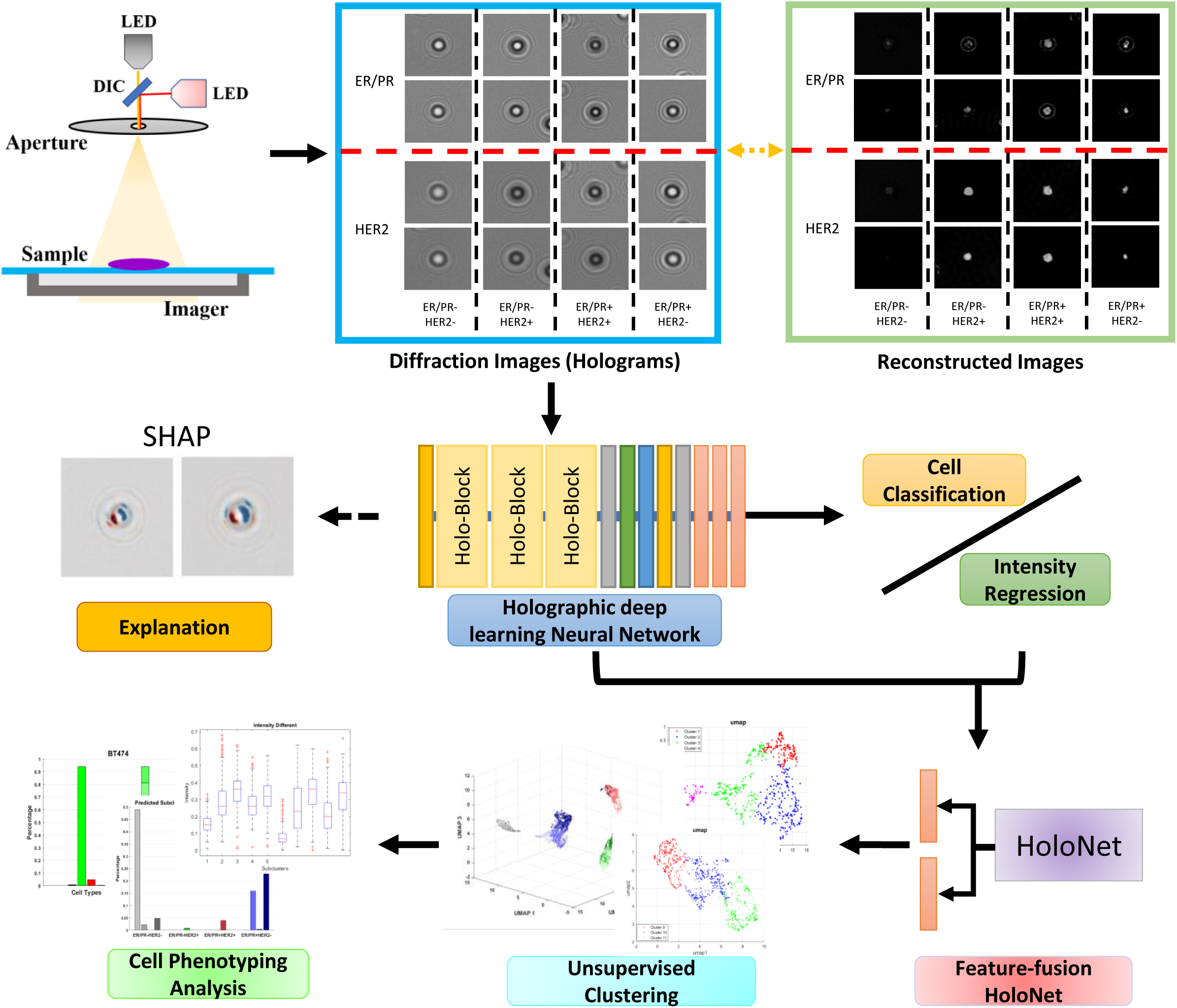
A workflow of the proposed deep learning models for cell phenotyping using lens-free digital in-line holography.

## Results

### Overview of Proposed Workflow

As described in **Figure 1**, the holograms were acquired by an LDIH imaging system from breast cancer cells that were color-stained for ER/PR and HER2. Specifically, cells were labeled with antibodies conjugated with oxidizing enzymes. Subsequent addition of chromogens generated visible color (red for ER/PR and blue for HER2), and then the stained cells were imaged with the LDIH system under multi-color illumination. We designed a novel holographic deep learning model firstly to classify four breast cancer cell types: ER/PR-HER2-, ER/PR-HER2+, ER/PR+HER2+, and ER/PR+HER2- and predict color intensities of ER/PR and HER2 staining. Second, we advanced our deep learning model to learn holographic features to identify subclusters hidden in heterogeneous breast cancer samples by combining manifold learning and unsupervised clustering.

### Hologram Classification and Regression by HoloNet

We design a novel deep learning HoloNet model to extract and analyze holographical features. **Figure 2(a)** shows the architecture of our proposed HoloNet model. A holo- block is built to combine local details of cellular objects with global features using a large kernel size of the convolutional filter and a concatenated layer. The HoloNet architecture is combined with a softmax layer as the output layer to classify breast cancer cell types. The cell classification results of the HoloNet model are shown in **Figure 2(b-c)**. Here we compare the classification performance with two types of input images, holograms, and reconstructed images. We also used CNN [33–35] and Resnet [36] models to compare the HoloNet model. First, we found that holograms provide more efficient features than reconstructed images in cell classification (**Figure 2(b)**). Even though the reconstructed images are built from holograms, detailed information may be lost during the numerical calculation of reconstruction processing. Overall, the HoloNet model with holograms performs better than other deep learning models trained with holograms or reconstructed images (**Figure 2(b)**). In addition, the HoloNet model achieved higher accuracy and F1- score than different DL approaches and achieved over 95.5% cell classification performance (**Figure 2(c); Supplementary Table 1**). HoloNet model improved the classification accuracy by 5% over the previous work using a standard CNN model [6]. HoloNet has a marginally better classification performance than DenseNet [37], which also combines multiple-scale image feature, However, it takes much less time to train HoloNet than DenseNet because it has much fewer model parameters (**Supplementary** Figure 1).

**Figure 2.**
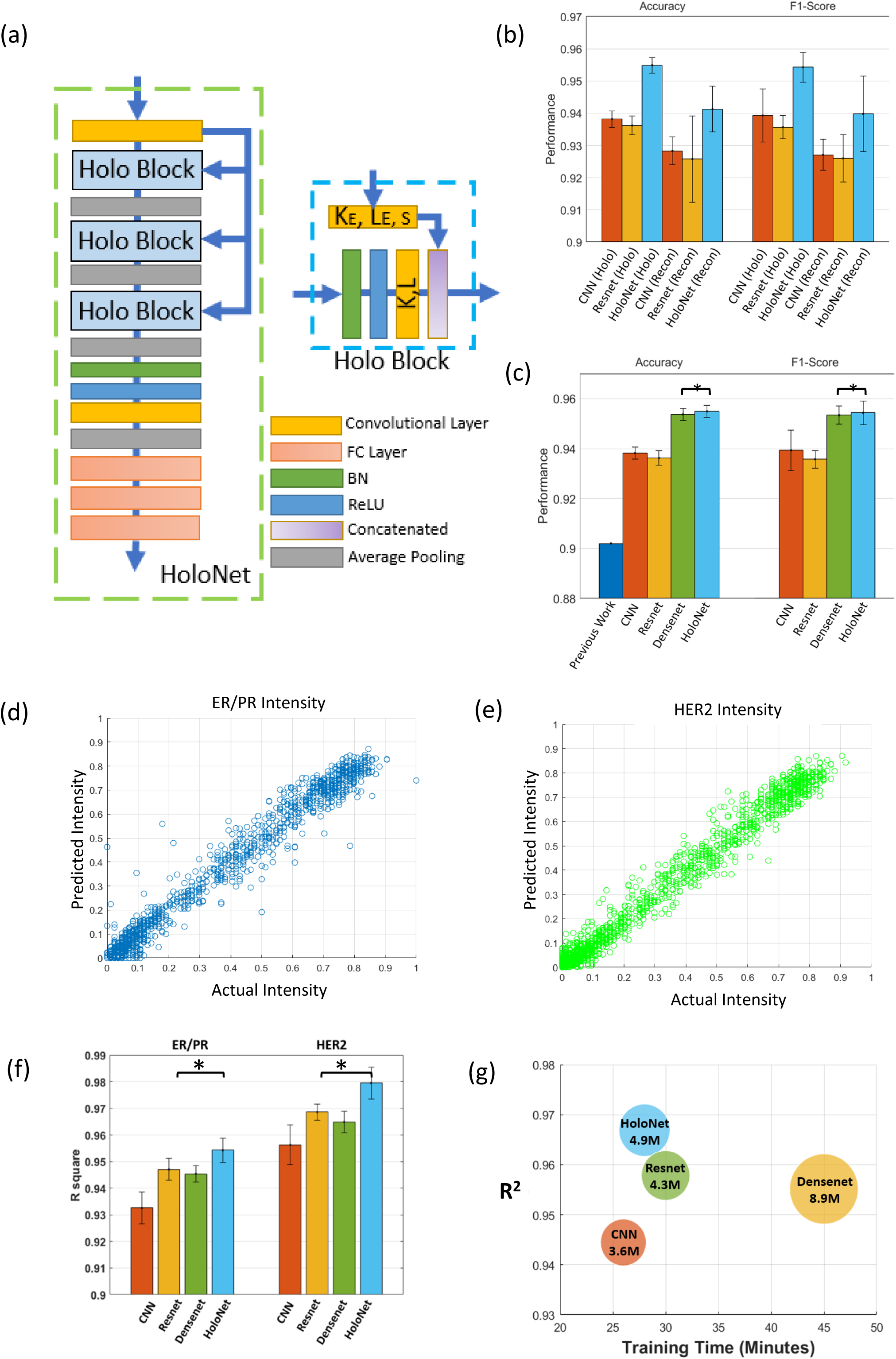
Structure and performance of the HoloNet. **(a)** The architecture of the HoloNet, where *K_E_*,*L_E_* and s represent the kernel size, number of feature layer, and sliding size, respectively. **(b-c)** Comparison of the performances of the classification of breast cancer cell phenotypes between holograms and reconstructed images (b), and among different deep learning structures (c). *p=4×10^-4^. The error bars represent the standard deviation. **(d-e)** Intensity regression results from the HoloNet. **(f)** Comparison of the ER/PR and HER2 regression performance among different deep learning structures. *p=6×10^-34^ (ER/PR), 3×10^-53^ (HER2). **(g)** Comparison of the regression performance and training times among different deep learning structures. The numbers in the circles are the numbers of the parameters in each deep learning model and the sizes of the circles are proportional to them.

Because the marker intensities of ER/PR or HER2 are the critical features for phenotyping breast cancer cells, we trained HoloNet with the fully connected layer of the regression output to quantify their intensities directly from raw holograms. The HoloNet regression model quantified the intensity values of ER/PR and HER2 channels without the reconstruction process, while the previous study did it using reconstructed images [6]. **Figure 2(d-e)** shows that the HoloNet model can accurately predict intensities in both ER/PR and HER2 staining channels, where the *R*^2^ scores are 0.9543 and 0.9795, respectively. We also trained other deep learning structures of CNN [33–35], ResNet [36], and DenseNet [37] to predict the intensity values, and compared their regression performance with that of HoloNet. As demonstrated in **Figure 2(f)** and **Supplementary** Figure 2, HoloNet significantly outperformed all of them in both ER/PR and HER2 channels. While HoloNet was slightly better for the classification task than DenseNet (**Figure 2(c)**), it significantly outperformed DenseNet for the regression task with a large margin. Also, the training of HoloNet is much faster than DenseNet or on par with CNN and ResNet (**Figure 2(g)**). Taken together, HoloNet is an accurate and efficient deep neural network for both classification and regression tasks with raw holograms.

To understand the superiority of HoloNet over CNN, ResNet, and DenseNet, we utilized SHAP (Shapely Additive exPlanations) values [38] to identify the hologram regions that play a significant role in enhancing classification and regression performance. The SHAP interpretation method, rooted in game theory principles, calculates SHAP values to measure the average marginal contribution of each feature [38]. These values provide a rigorous and comprehensive understanding of the importance and influence of individual features on the predictions made by deep learning models. By quantifying the contribution of each feature, SHAP values enable a more accurate interpretation of the model’s predictions.

The SHAP value distributions in **Figure 3(a)-(d)** indicate positive (red) or negative (blue) contributions to the cell type classification with different deep learning models. The SHAP values of HoloNet exhibit significantly less noise compared to other DL models. Additionally, HoloNet demonstrates a greater emphasis on specific and well-defined regions within diffraction patterns when compared to those other models. Consequently, HoloNet offers more precise and robust cell type classification capabilities. In the context of intensity regression (**Figure 3(e)**), the SHAP values of CNN, ResNet, and DenseNet demonstrate reduced noise compared to the classification task. However, these models encountered challenges in accurately identifying the specific hologram structures that were successfully recognized by HoloNet. Notably, in the cell type ER/PR-HER2-, which exhibits low intensity in both the 470 and 625 nm channels, HoloNet’s SHAP values reveal highly distinct regions. These distinct regions are particularly valuable for making accurate predictions in cases involving low intensity.

**Figure 3.**
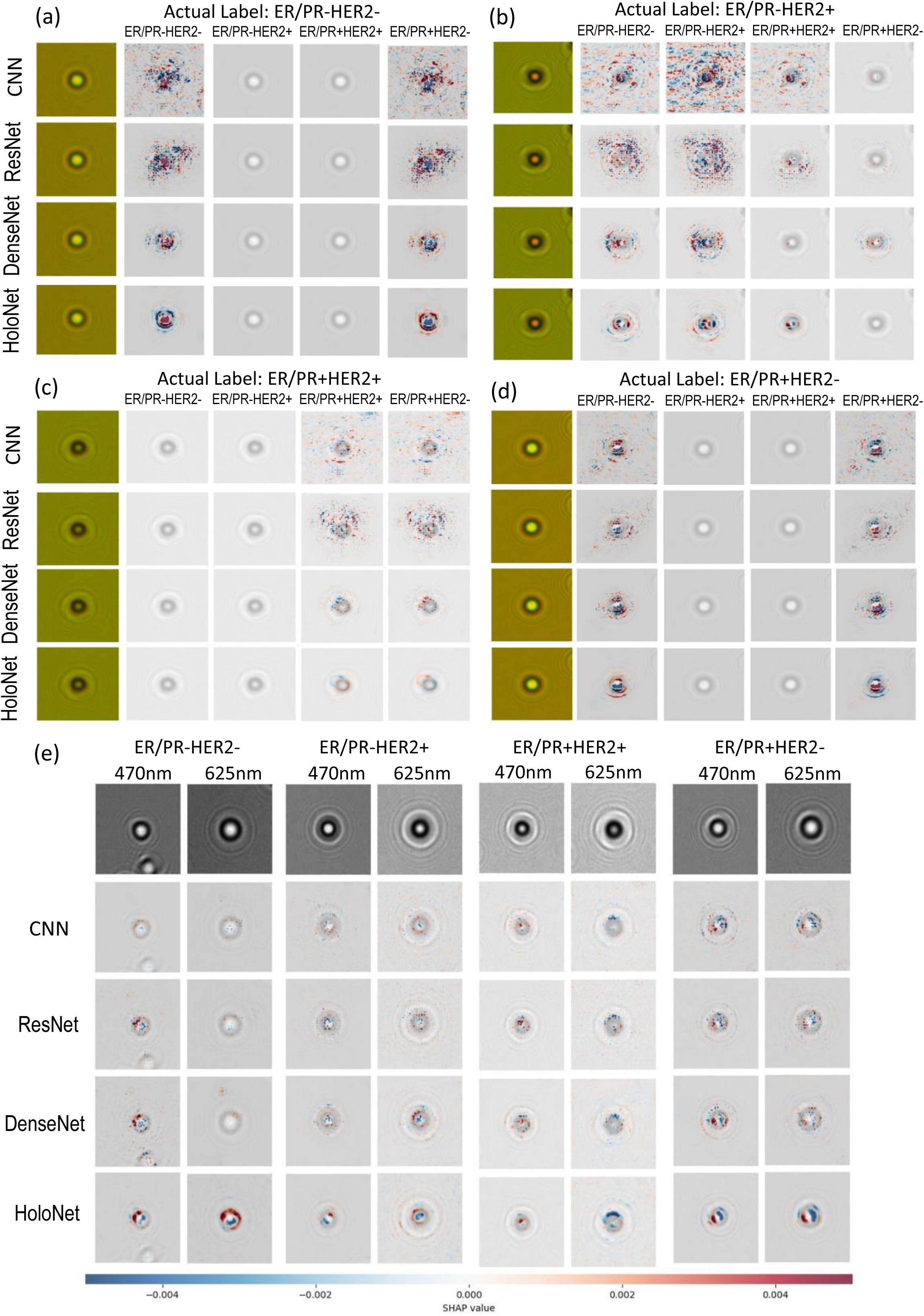
The Interpretability analysis of the HoloNet model. (a-e) Comparison of the SHAP value distributions on holograms for cell type classification (a-d) and marker intensity regression in 470 and 625 nm channels (e) among CNN, ResNet, DenseNet, and HoloNet.

### Sub-clustering of Breast Cancer Cells with Feature-Fusion HoloNet

Since deep learning models can extract rich features from input images, we advanced hologram feature embedding methods to identify subtypes of breast cancer cells. **Figure 4(a)** shows the workflow of feature extraction and sub-clustering analysis. We developed the feature-fusion HoloNet model to obtain the diffraction feature vectors. The hologram features extracted from HoloNet were used for the cell type classification and the intensity regression simultaneously. By performing multi-task learning, the HoloNet fused the regression features with the classification features to pay more attention to the influence of the features related to the intensities. Therefore, the resulting subclusters can have differential intensity distributions in each cell type. Then the feature vector is obtained from the feature-fusion HoloNet model and processed by Uniform Manifold Approximation and Projection (UMAP) method [39] to learn the feature manifold and reduce the dimension of features. **Figure 4(b)** and **4(c)** show the feature distributions of breast cancer cells obtained by the HoloNet with holographic input images and reconstructed input images, respectively. The feature distribution map from the reconstructed input images shows that the clusters of different cell types can be distinctly separated because of the distance. But, due to the lack of intra-cluster variability, it is difficult to find potential sub-clusters. The feature distribution map from holograms exhibited greater intra-cluster variability while the inter-cluster distances became smaller than the features from the reconstruction images. In contrast, the hologram distribution map from the feature-fusion HoloNet model in **Figure 4(d)** showed the sizable intra-cluster variability and inter-cluster distances, suggesting that the features from the feature-fusion HoloNet model are suitable for sub-clustering analysis.

**Figure 4.**
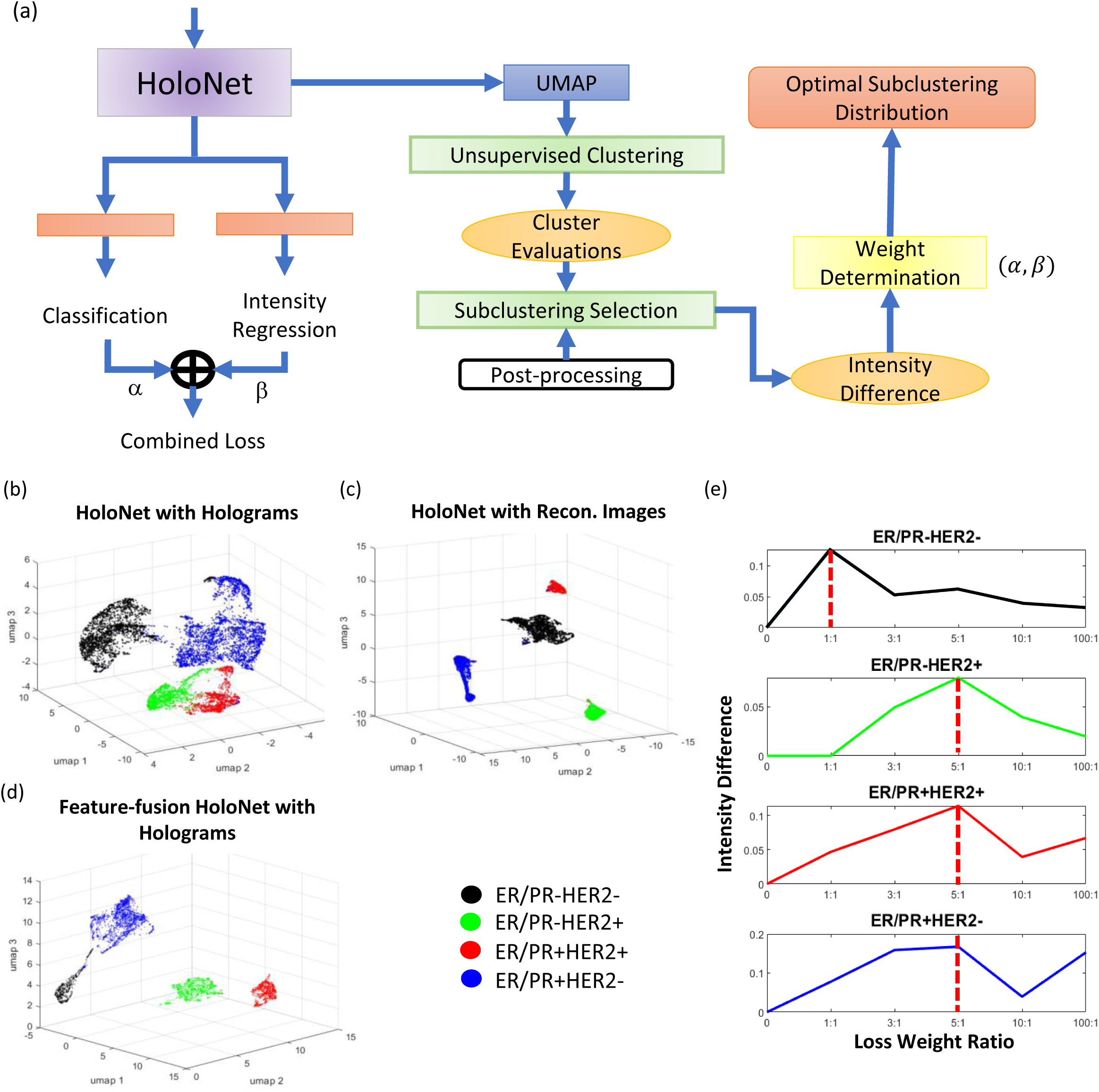
Pipeline of HoloNet-based unsupervised learning. (a) The architecture of the feature-fusion HoloNet with unsupervised subclustering. (**b-d**) Feature distribution maps from the HoloNet with holograms (b), the HoloNet with reconstructed images (c). the feature-fusion HoloNet with holograms (d). (**e**) Determination of the loss weight in each phenotype of breast cancer cells.

To determine the optimal number of subclusters in each cell type, we combined community detection with spectral clustering [40] and grid search. We used clustering evaluation functions to estimate the cohesion values of subclusters and used rank voting to select the optimal number of subclusters in each class. To determine the optimal balance between the branches of classification and regression, we varied the ratio of two loss functions, and evaluated the sub-clustering results by the mean intensity differences of ER/PR and HER2 among the subclusters. **Figure 4(e)** shows the mean intensity values in different ratios of the loss of classification and regression, and we chose the embedding that provided the maximum mean intensity differences.

### Identifying Subclusters of Breast Cancer Cells

Figure 5(a) represents the subcluster distribution maps with the optimal loss weight in each cell type. We obtained four subclusters with loss weight ratio 1:1 in ER/PR-HER2-, four subclusters with loss weight ratio 5:1 in ER/PR-HER2+ and ER/PR+HER2- cell types, and three subclusters with loss weight ratio 5:1 in ER/PR+HER2+. After the subclusters were obtained in different cell type groups, we observed the cell population of subclusters in cell line sample cases. In Figure 5(b), we found that MCF7 and T47D cell lines are dominated by ER/PR+HER2- cell type, but they consist of markedly different distributions of the subclusters. While Cluster 13 is the major subcluster in MCF7, Cluster 15 is the major component in T47D. Moreover, the SKBR3 cell line whose primary cell type is ER/PR+HER2+ consists of Cluster 9-11 equally. In the BT474 cell line, Cluster 6 and 7 dominate the ER/PR-HER2+ cell type. MDA-MD-231 cell line where most of the cells belong to ER/PR-HER2- mainly composed Cluster 1-3. In addition, Cluster 3 and 4 exist in ER/PR-HER2- of the MCF7 cell line, and Cluster 8 mostly dominates the ER/PR- HER2+ cell type of the SKBR3 cell line.

**Figure 5.**
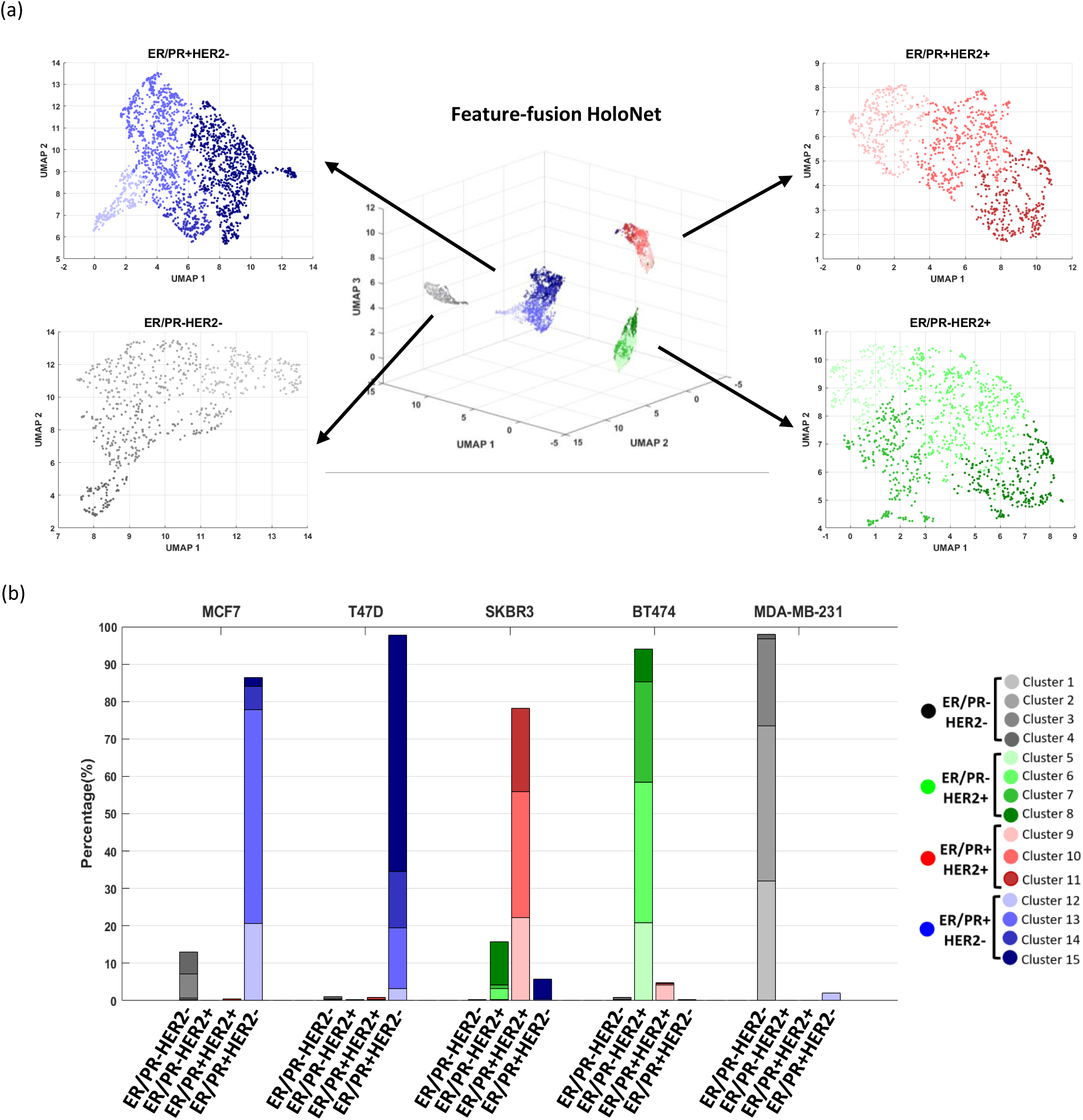
Subcluster results from hologram embedding. (**a**) Distribution of hologram features from the feature-fusion HoloNet embedding and the subclustering results in each cell type. (**b**) Subcluster distribution in each phenotype of breast cancer cells in each breast cancer cell line.

To know the natures of these subclusters, we quantified the average marker intensities in each subcluster after spectral unmixing of the marker channels (see Methods) [6] (Figure 6(a)**-(d), Supplementary** Figure 3(a)**-(d)** for the intensities before the spectral unmixing) and the distributions of breast cancer cell features in each subcluster (Figure 6(e)**-(l)** for UMAP distribution**, Supplementary** Figure 3(e)**-(h)** for intensity distribution). Our analyses revealed the existence of the overlooked subclusters of cancer cell phenotypes that exhibit the characteristics of multiple breast cancer phenotypes. Because we used the feature-fusion HoloNet model to learn the features partially discriminative to marker intensities, the mean intensity values of the subclusters were generally statistically different (Figure 6(a)**-(d), Supplementary** Figure 3(a)**-(d)** for the intensities before the spectral unmixing). In ER/PR-HER2- cell type, the mean intensity values of Cluster 1 and 2 in both markers are the lowest while those of Cluster 4 are the highest (Figure 6(a)). The mean ER/PR intensities of Cluster 3 are medium and significantly higher than those of Cluster 1 and 2. (Figure 6(a)). Most of the cells in Cluster 1 and 2 were from MDA-MB-231. Cluster 3 has a mixed population of MDA-MB-231 and MCF7. Cluster 4 mainly consists of MCF7 along with minor populations from T47D, BT474, and MDA-MB-231 (Figure 6(e) and **(i)**). In ER/PR-HER2+ cell type, the subclusters also had different mean marker intensity values (Figure 6(b)). Cluster 5, 6, and 7 were mainly from BT474. The mean HER2 intensity value in Cluster 7 is the highest among the subclusters, and Cluster 8, in which BT474 and SKBR3 co-existed equally (Figure 6(f) and **(j)**), has the highest ER/PR intensity value. In ER/PR+HER2+ cell type, the mean ER/PR intensity values increased from Cluster 9 to 11 while the mean HER2 intensity values decreased (Figure 6(c)). The primary cell line in these subclusters is SKBR3, but Cluster 9 has a minor cell population from BT474 (Figure 6(g) and **(k)**). In ER/PR+HER2- cell type, the mean intensity values of both markers tended to increase from Cluster 12 to 15 (Figure 6(d)). Cluster 12 and 13 mainly consisted of MCF7 along with minor proportions of T47D. Cluster 14 and 15 mainly consisted of T47D along with minor proportions of MCF7 in Cluster 14 and SKBR3 in Cluster 15 (Figure 6(h) and **(l)**).

**Figure 6.**
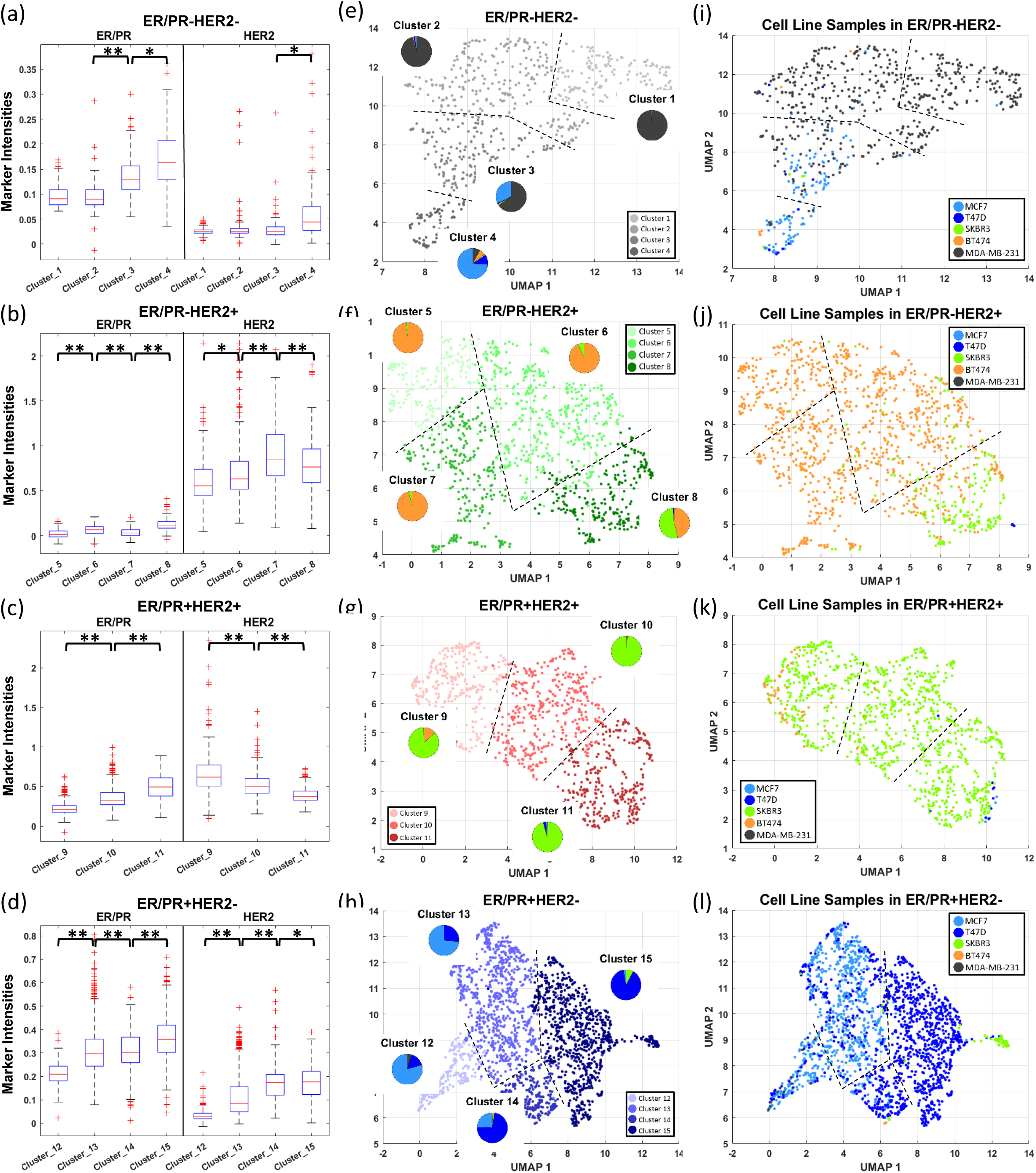
Characteristics of the identified subclusters of breast cancer cells. (**a-d**) Differences of the mean intensity values of the subclusters in each breast cancer cell type. (**e-h**) UMAP visualization of hologram features color-coded with the subclusters. The pie plots indicate the proportion of the cell lines in each subcluster (the color code of cell lines are in (i-l)). (**i-l**) UMAP visualization of hologram features color-coded with the cell lines. *p=10^-10^∼10^-4^ and **p<10^-10^ . Box plot displays the minimum, first quartile, median, third quartile, maximum, interquartile range, whiskers, and outliers.

As summarized in Table 1, in Cluster 3, 4, 8, 9, 14, and 15, the cells from different cell types co-exist. Cluster 3 is near the phenotypic boundary between ER/PR-HER2- and ER/PR+HER2-. Cluster 4 is the boundary among ER/PR-HER2-, ER/PR-HER2+ and ER/PR+HER2-. Cluster 8 and 9 are the boundary between ER/PR-HER2+ and ER/PR+HER2-. Cluster 14 and 15, consisting of T47D are the boundary between ER/PR+HER2- and ER/PR+HER2+. While some breast cancer cells exhibit the characteristics shared by multiple cell phenotypes, they were considered to be in a single cell phenotype for a diagnostic purpose. Here, we were able to identify those overlooked phenotypes of breast cancer cells using the feature-fusion HoloNet.

**Table 1.** Summary of the distributions of breast cancer cell lines in the identified subclusters.

| Cell Type | ER/PR-HER2- |  |  |  | ER/PR-HER2+ |  |  |  | ER/PR+HER2+ |  |  | ER/PR+HER2- |  |  |  |
| --- | --- | --- | --- | --- | --- | --- | --- | --- | --- | --- | --- | --- | --- | --- | --- |
| Subcluster | 1 | 2 | 3 | 4 | 5 | 6 | 7 | 8 | 9 | 10 | 11 | 12 | 13 | 14 | 15 |
| MCF7 |  |  | + | ++ |  |  |  |  |  |  |  | ++ | ++ | + |  |
| T47D |  |  |  | + |  |  |  |  |  |  |  | + | + | ++ | ++ |
| SKBR3 |  |  |  |  |  |  |  | + | ++ | ++ | ++ |  |  |  | + |
| BT474 |  |  |  | + | ++ | ++ | ++ | + | + |  |  |  |  |  |  |
| MDA-MB-231 | ++ | ++ | ++ | + |  |  |  |  |  |  |  |  |  |  |  |

We also performed the same sub-clustering using reconstructed images to demonstrate the effectiveness of the raw hologram features. We quantified the average marker intensities in each subcluster (**Supplementary** Figure 4(a)**-(d), Supplementary** Figure 5(a)**-(d)**) for the intensities before the spectral unmixing) and the distributions of breast cancer cell features in each subcluster (**Supplementary** Figure 4(e)**-(l)** for UMAP distribution**, Supplementary** Figure 5(e)**-(h)** for intensity distribution). Even though there exists cell line specificity in the feature distribution (**Supplementary** Figure 4**(i-l)**), the cell lines in ER/PR-HER2- (**Supplementary** Figure 4**(e-i)**) and ER/PR-HER2+ (**Supplementary** Figure 4**(h-l)**) are not well clustered like the hologram cases. MCF7 cells are less enriched in Cluster 5 from the reconstructed images in ER/PR-HER2- than Cluster 4 from the hologram (Figure 6(e) and **Supplementary** Figure 4(e)). In ER/PR+HER2-, the sub-clustering of the holograms was able to distinguish MCF7 and MDA-MB-231 clearly (Figure 6(h)), whereas sub-clustering of the reconstructed images could not (**Supplementary** Figure 4(h)). In the cases of ER/PR-HER2+ and ER/PR+HER2+, the sub-clustering of the reconstructed images was able to isolate the subclusters specific to the cell lines, SKBR3 in Cluster 7 (**Supplementary** Figure 4**(f-j)**) and BT474 in Cluster 8 (**Supplementary** Figure 4**(g-k)**), consistent with the results from the holograms (Figure 6**(f-j)** and **6(g-k)**). These results suggest that raw holograms provide the features that allow more distinct sub-clustering than reconstructed images.

To compare the features learned by HoloNet and DenseNet from raw holograms, we replaced HoloNet with DenseNet in our feature-fusion pipeline. First of all, the training curve showed that HoloNet-based feature-fusion model had lower training and validation losses than DenseNet-based feature fusion model (**Supplementary** Figure 6). In ER/PR- HER2- and ER/PR+HER2-, where two major cell lines exist in similar proportions, DenseNet showed a similar capability to divide these cell lines in comparison to HoloNet (**Supplementary** Figure 7(i) vs Figure 6(i), **Supplementary** Figure 7(l) vs Figure 6(l)). In contrast, in ER/PR-HER2+ and ER/PR+HER2+ which are dominated by major cell lines, HoloNet isolate minor cell lines better than DenseNet: SKBR3 cells in ER/PR-HER+ were more spread on the UMAP of DenseNet than HoloNet (**Supplementary** Figure 7(j) vs Figure 6(j)). Also, while T47D and MCF7 were located on the opposite side of the UMAP from HoloNet, they coexisted on the same side of the UMAP from DenseNet. This suggests that HoloNet produced the features that are more suitable for identifying minor cell populations.

### Existence of Discovered Subclusters in Patient Samples

Our subclustering analysis was conducted on multiple breast cancer cell lines to explore the clinical relevance of these subclusters. To validate their significance in an actual clinical context, we utilized hologram data from two breast cancer patient cases [6]. We employed the proposed HoloNet classification model to determine the distribution of cell types within these patient cases. The proportions of cell types for the two breast cancer patients are illustrated in Figure 7(a). Although the most prominent cell types are ER/PR- HER2- in Patient #1 and ER/PR+HER2- in Patient #2, the overall distributions of cell types show only marginal differences. By utilizing our feature-fusion HoloNet model, we generated feature vectors of cellular holograms from these patients and matched them against our subclusters of breast cancer cells. The results conclusively confirmed the existence of subclusters within the patient samples. In Figure 7(b), we found that Patient #1 has much more Cluster 1 from ER/PR-HER2- than Patient # 2 while Patient #2 has much more Cluster 4 from ER/PR-HER2- than Patient case 1. Given our finding that Cluster 4 share the phenotypes from ER/PR-HER2+ and ER/PR+HER2-. The characteristics of ER/PR-HER2- in these two patients are considered to be distinct.

**Figure 7.**
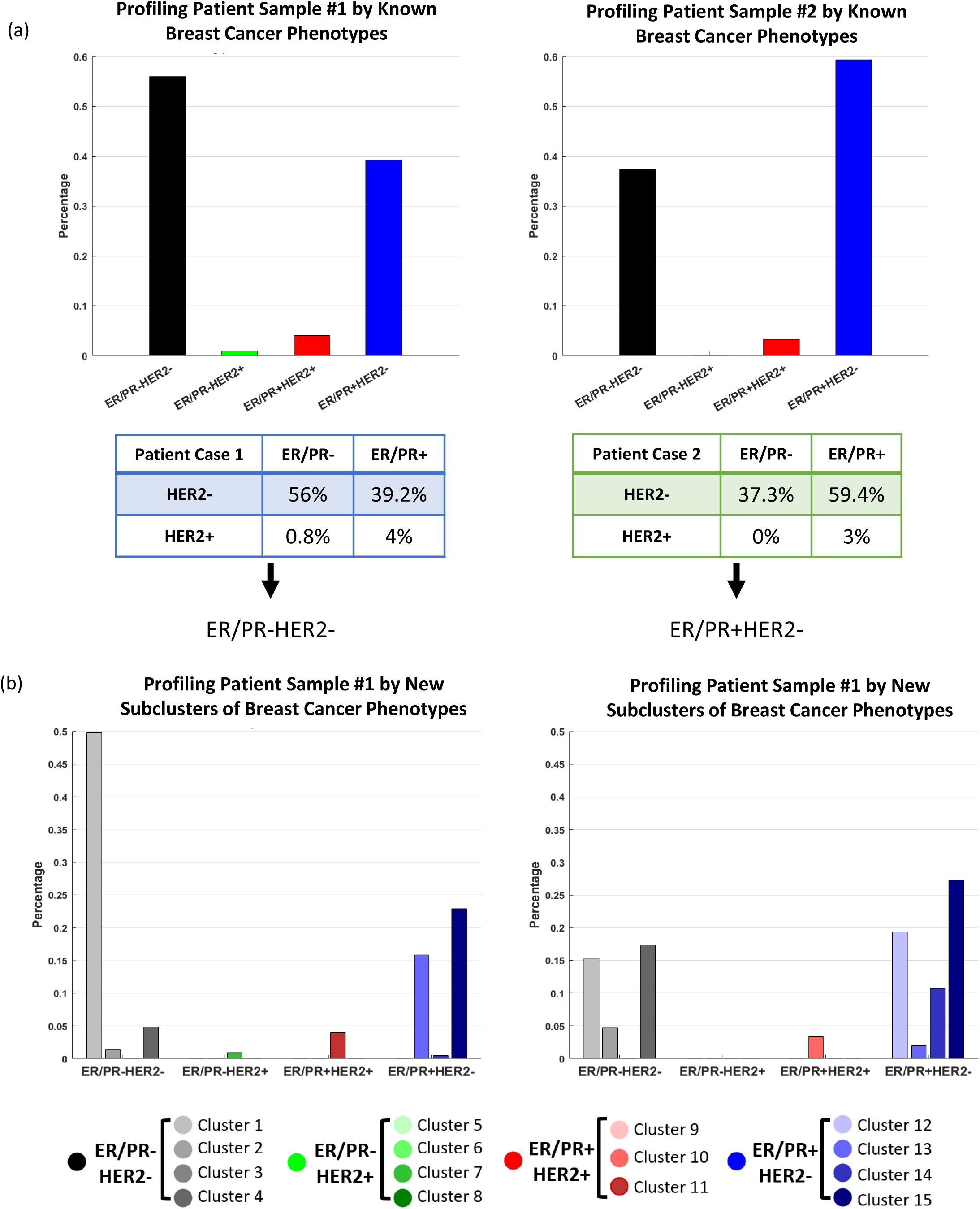
Existence of breast cancer subclusters in patient samples. (**a**) Profiling two patients using the known phenotypes of breast cancer cells. (**b**) Profiling two patients using the discovered subclusters of phenotypes of breast cancer cells

For ER/PR+HER2- type, two patients have the similar most considerable proportions of Cluster 15, sharing the similar characteristics of ER/PR+HER2+. However, Patient #1 has a much more proportion of Cluster 13 than Patient # 2, while Patient #2 has much more Cluster 12. Since there are significant differences in both channels between Cluster 12 and 13, the cellular phenotypes in ER/PR+HER2- of these patients are also distinct. Taken together, we demonstrated that some of the identified subclusters existed in the breast cancer patient samples, and the sub-clustering results could provide much richer information on the disease status.

## Discussion

We developed the HoloNet, which can efficiently learn high-level diffraction features from raw holograms to precisely discriminate breast cancer cell types in both supervised and unsupervised learning settings. Significantly, the holo-block unit adapts different large-scale filters to collect multi-scale cell feature information to identify detailed local cellular features. It is because the local feature information in holograms does not correspond to a particular part of cellular images but rather the entire images. The large-scale filters applied to holograms can collect more related local cellular information. Our HoloNet can efficiently extract cell information from holograms to provide better performances of cell classification and intensity regression than other existing deep learning models, in addition to superior interpretability.

We demonstrated that the feature embedding directly from holograms enabled us to identify detailed subclusters of breast cancer cells. We added the intensity regression into the feature-fusion HoloNet model together with the classification of the previously known cell types. This structure helped the neural network pay more attention to specific hologram features to enhance the difference of marker intensities among different cell types. Then we optimized the loss weights to maximize the differences of the marker intensities among potential subclusters. This hologram embedding allowed us to identify the subclusters within the known cell types for refined cellular phenotyping. Some of the identified subclusters have the phenotypes shared by multiple breast cancer cell types since they are located near the class boundaries in the feature space. These rare and subtle cellular phenotypes may have implications in drug sensitivity and resistance, which should be further validated in future studies. We expect that HoloNet, in conjunction with LDIH, opens a new opportunity to fully characterize intra-tumor and inter-tumor heterogeneity in breast cancer and provide clinicians with valuable information for patient-specific breast cancer therapy.

## Methods

### Data collection

Breast cancer cells were captured by the surface coated by antibodies (HER2, EpCAM, EGFR, and MUC1) [6] and bio-adhesives, and then stained by anti-ER/PR and anti-HER2 conjugated with chromogen. Then cellular holograms were obtained by an LDIH system, and the image patches containing a cell were cropped with 64Х64 pixel size in both hormonal staining channels. These image data include four different cell types: ER/PR- HER2-, ER/PR-HER2+, ER/PR+HER2-, and ER/PR+HER2+, from five cell lines (MCF7, T47D, SKBR3, BT474, and MDA-MB-231) [6]. The number of all holograms was 5026. For more efficient training, data augmentation was applied to balance the size of different cell types by using random rotation and flipping. Then we split the data into training (80%), and testing (20%) with 5-fold cross-validation for model evaluation. In each fold, the validation set is 18.75% of the training set (15% of the whole data). The ground truth of intensity values of ER/PR and HER2 of cell images were obtained from the reconstructed images.

### Holographical deep learning network (HoloNet)

We constructed the HoloNet based on the convolutional neural networks by adding several large-kernel-size filters to efficiently extract hologram features. Here we designed a block called holo-block, which combines local and global holographical features. The parameters K, L, and s in a holo-block (Figure 2(a)) represent the kernel size of the convolutional layer, number of feature layers, and sliding, respectively. There are three holo-blocks with different kernel sizes of convolutional layers, which are 16X16, 24X24, and 32X32, respectively, and average pooling layers are used to obtain and emphasize specific feature information. The number of feature layers in these blocks was 64. The sliding numbers were set to 1, 2, and 4 for each holo-block. Moreover, a convolutional layer with batch normalization and three fully connected layers are connected to these holo-blocks to build the HoloNet model. Rectified linear unit (ReLu) was used as an activation function in the model [41]. Based on the HoloNet architecture, we implemented a DL model to classify four types of breast cancer cells by a softmax layer. Then, for the intensity regression, we constructed the HoloNet architecture with a fully connected layer to predict intensity values of ER/PR and HER2 staining channels from holograms.

### Feature-fusion HoloNet

The architecture of the feature-fusion HoloNet model includes the HoloNet model with two fully connected layers of intensity regression and the cell type classification (Figure 3(a)). The total loss function of the training is shown as:

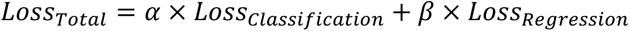

where α and β are the loss weights of classification and intensity regression for loss balancing. Here we used Brier loss [42] for the classification loss as below:

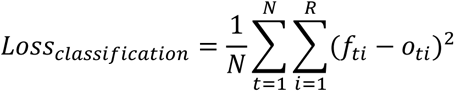

where N is the number of observations and R means the number of categorical labels. *f* and *o* represent the predictive and true label distribution, respectively. Brier loss function is similar to mean square error loss but has the same ability of loss energy as the cross-entropy function.

### Neural Network Training

We used Adam optimizer with learning rate=10^−4^ for classification, stochastic gradient decent optimizer with moment=0.9 for regression, and batch size=128. The categorical cross-entropy was used as a loss function for training the HoloNet model for cell classification, and the mean square error loss function was used for HoloNet model training for the intensity regression. We set the input image size as 64×64×2 for the classification and 64×64 in each staining channel for the regression. For the HoloNet model, the maximum epoch was 100 for the classification and 500 for the regression. The pixel value of input image was normalized from 0 to 1. We also automatically reduce the learning rate by multiplying with 0.1 in every 20 epochs. For training the feature-fusion HoloNet model, we set the brier loss function as a loss function for the classification and the maximum epoch=150. We used default parameters in the Keras library, and the environment in Python was TensorFlow 1.15 with CUDA 10.0 for both HoloNet models.

### Unsupervised Clustering and Subcluster Selection

We used the second-to-last layer of the feature-fusion HoloNet model to extract the feature vector of 500 dimensions, and the UMAP method [39] was used to reduce these 500 dimensional feature vectors to three-dimensional space. In the parameters of the UMAP method, the number of the neighborhood was set to 20 and the dimension of the space was 3. Also, the minimum distance among the observation was set to 0.1, and the string metric was the correlation to compute distance in high dimensional space. Then, the pairwise distances between hologram feature vectors were calculated to generate a similarity matrix. The threshold value of similarity was set to 0.5 to build an adjacent matrix. This adjacent matrix was used as community detection to obtain subclusters by using spectral clustering [40]. The grid search method was used to find the optimal number of subclusters in each cell type. We set the number of subclusters from 1 to 10 and evaluated the clustering quality of different clustering numbers using clustering evaluation functions including silhouette coefficient [43], Dunn’s index [44], Calinski-Harabasz index [45], and Davies-Bouldin index [46]. Then the optimal number of subclusters in each cell type was selected by voting the highest rank from the list of clustering values among different subclustering numbers.

### Spectral Unmixing of Chromogens

There exists an overlap of the absorption spectra between the chromogens used in this study. Therefore, spectral unmixing, which determine the contributions from each chromogen, was performed according to the previous work [6] as follows.

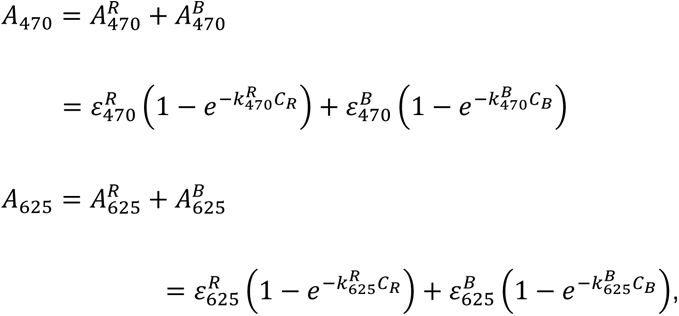

where *A*_470_ and *A*_625_ represent the measured intensities of ER/PR and HER2, respectively, and *C_R_* and *C_B_* represent the actual contributions of the chromogens. The parameters *ε*_470_*^R^*, *ε*_470_*^B^*, *ε*_625_*^R^* and *ε*_625_*^B^* were experimentally evaluated to be 0.806, 0.2812, 0.3352, and 0.6517. Also, *k*_470_*^R^*, *k*_470_*^B^*, *k*_625_*^R^* and *k*_625_*^B^* were to be 1.3345, 2.3985, 2.346, and 2.848. *C_R_* and *C_B_* are obtained by solving the system of these nonlinear equations.

### Loss Weight Optimization

We used grid search to determine the optimal ratio of loss weight in each cell type for subclustering. We calculated Euclidean distances of the mean intensity values of ER/PR and HER2 channels among subclusters and then averaged those Euclidean distances. We evaluated this mean Euclidean intensity distance among the subclusters with the varying loss ratios: 1:1, 3:1, 5:1, 10:1 and 100:1 (classification : regression). Then we select the optimal weight combinations which provide the maximum intensity difference in each cell type (Figure 3(e)).

### Quantification and Statistical Analysis

For Figure 2(b), the accuracy and F1 score are shown as the mean with standard errors. Here the two-sided Wilcoxon rank sum test is used to test statistical significance between the model performances without the assumption of the normal distribution of the data. The statistical test is considered significant if the p-value is less than 0.01. For Figure 2(d), the model performance was presented using R^2^ values of linear regression. The squares of residuals were used to calculate p-values. The two-side Wilcoxon rank sum test was used to test statistical significance without the assumption of the normal distribution of the data. The statistical test is considered significant if the p-value is less than 0.01. For Figure 6**(a-d)** and Figure 7**(a-d)**, the two-side unpaired Wilcoxon rank sum test was used to test statistical significance between the marker intensity distributions of two subclusters without the assumption of the normal distribution of the data. If the p-value is less than 0.01, the statistical test is considered significant. The quantification and statistical analyses are performed in MATLAB (2020b).

### Model Interpretation

To analyze and gain insights into the distribution of crucial features in various deep learning models, we employed the GradientExplainer module from the SHAP python library. This Explainer module enables the identification of feature contributions in both cell classification and intensity regression tasks by computing the gradients of the target feature function as SHAP values. For the classification task, we utilized pre-trained models such as CNN, ResNet, DenseNet, and HoloNet to generate SHAP values for different classes within the test dataset. Subsequently, we visualized the positive and negative influences of feature distributions for each class identification. In the intensity regression task, the SHAP values also provided valuable insights into the specific impact of features on predicting intensity values.

## Data availability statement

The datasets used in the current study are available from the corresponding author on a reasonable request.

## Code availability statement

The code used in the current study is available from the corresponding author on a reasonable request.

## Acknowledgement

We thank Boston Scientific for providing us with the gift for deep learning research. This work was supported by Department of Defense, United States (Grant Numbers: W81XWH1910200 for K.L. and W81XWH1910199 for H.L.)

## Competing Interests

The author declares no competing financial or non-financial interests.

### Author Contributions

T.S. initiated the project, designed the pipeline, and wrote the final version of the manuscript and supplement. M.C. wrote the code for the regression model. J.M. and H.I. set up the imaging system and generated the hologram data. K.L. and H.L. coordinated the study and wrote the final version of the manuscript and supplement. All authors discussed the results of the study.

## Author Information

Correspondence and requests for materials, data, and code should be addressed to K.L. or H.L.

---

**Supplementary Figure 1.**
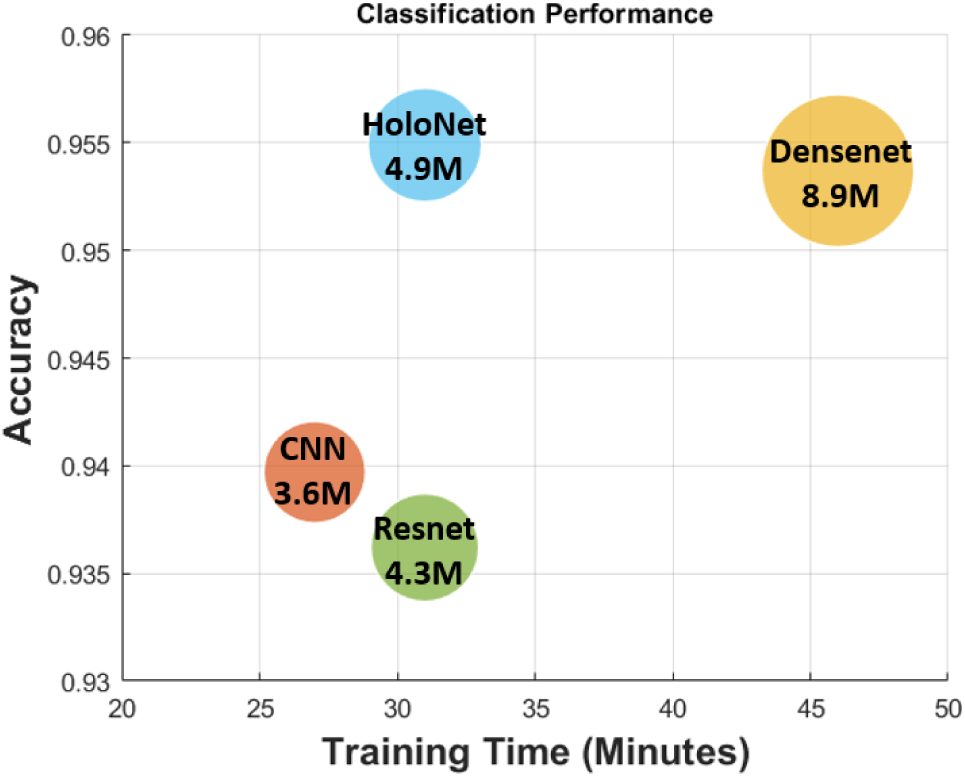
Comparison of the classification performance and training times among different deep learning structures. The numbers in the circles are the numbers of the parameters in each deep learning model and the sizes of the circles are proportional to them.

**Supplementary Figure 2.**
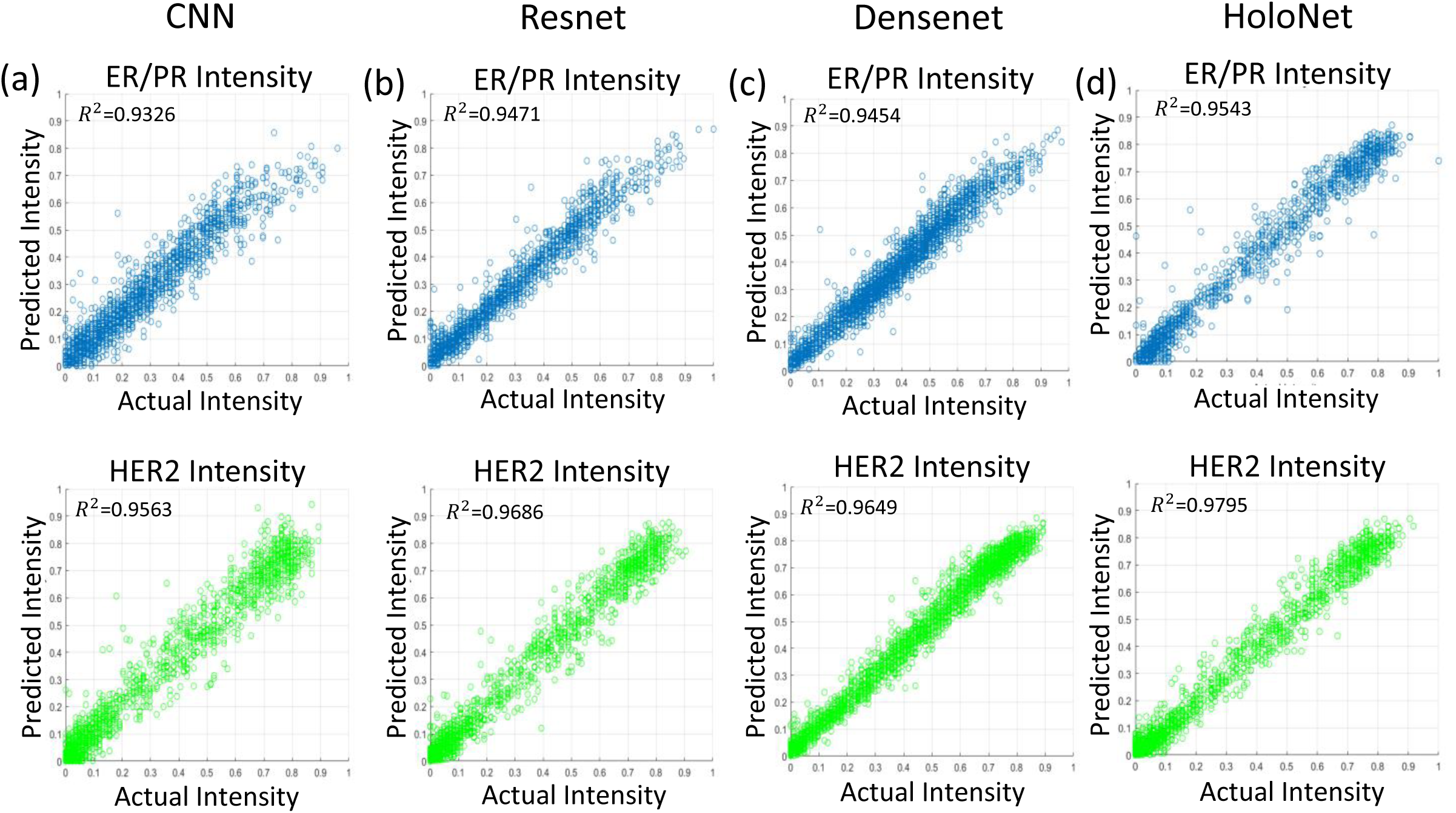
The ER/PR and HER2 intensity regression results of (a) CNN, (b) Resnet, (c) DenseNet, and (d) HoloNet models are shown. The proposed HoloNet model provides better regression performance than others in both staining channels.

**Supplementary Figure 3.**
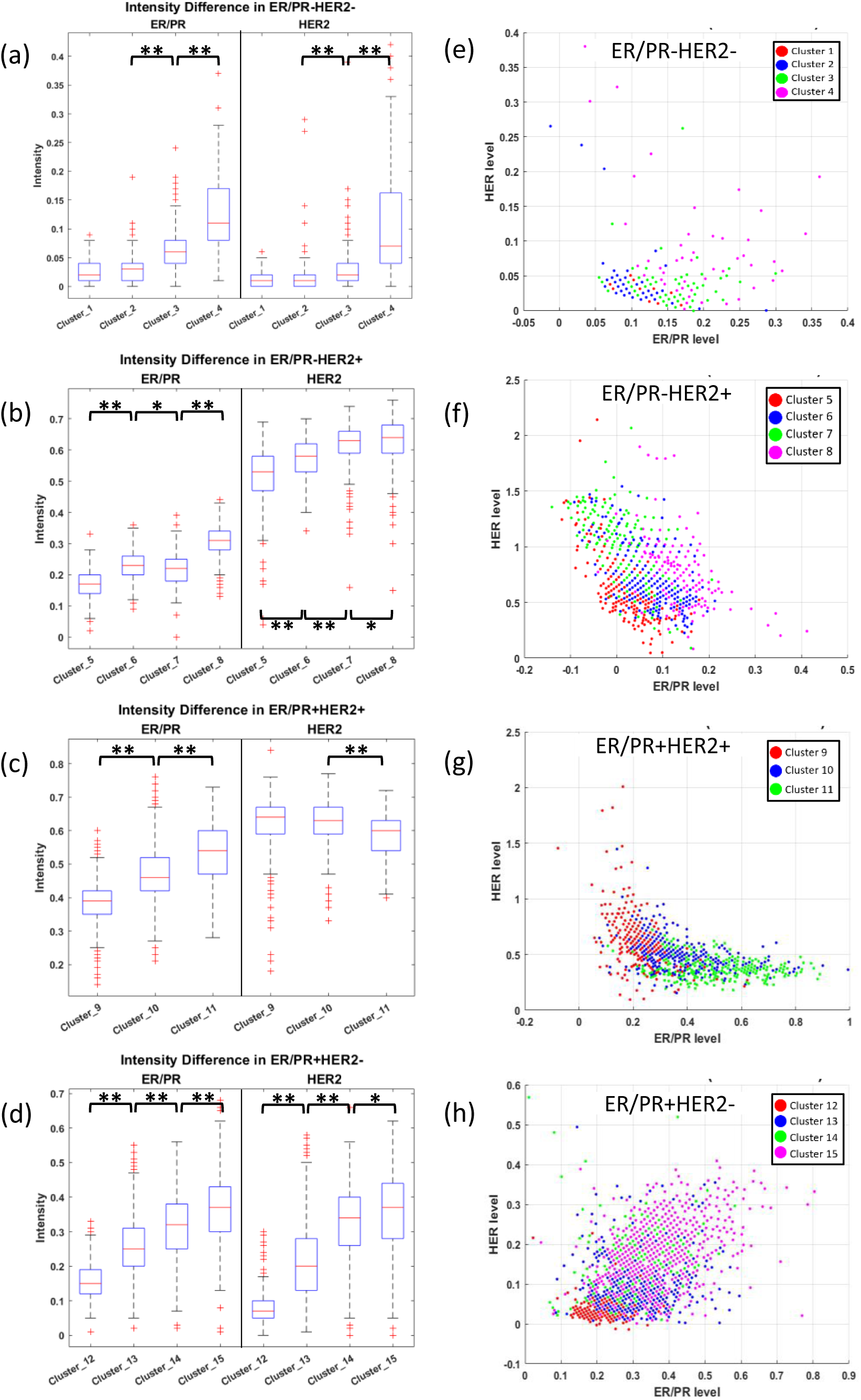
Characteristics of the identified subclusters of breast cancer cells. **(a-d)** Differences of the mean raw intensity values of the subclusters in each breast cancer cell type. **(e-h)** Distribution of marker intensities in each breast cancer cells color-coded with the subclusters. *p=10^-10^∼10^-2^ and **p<10^-10^ .

**Supplementary Figure 4.**
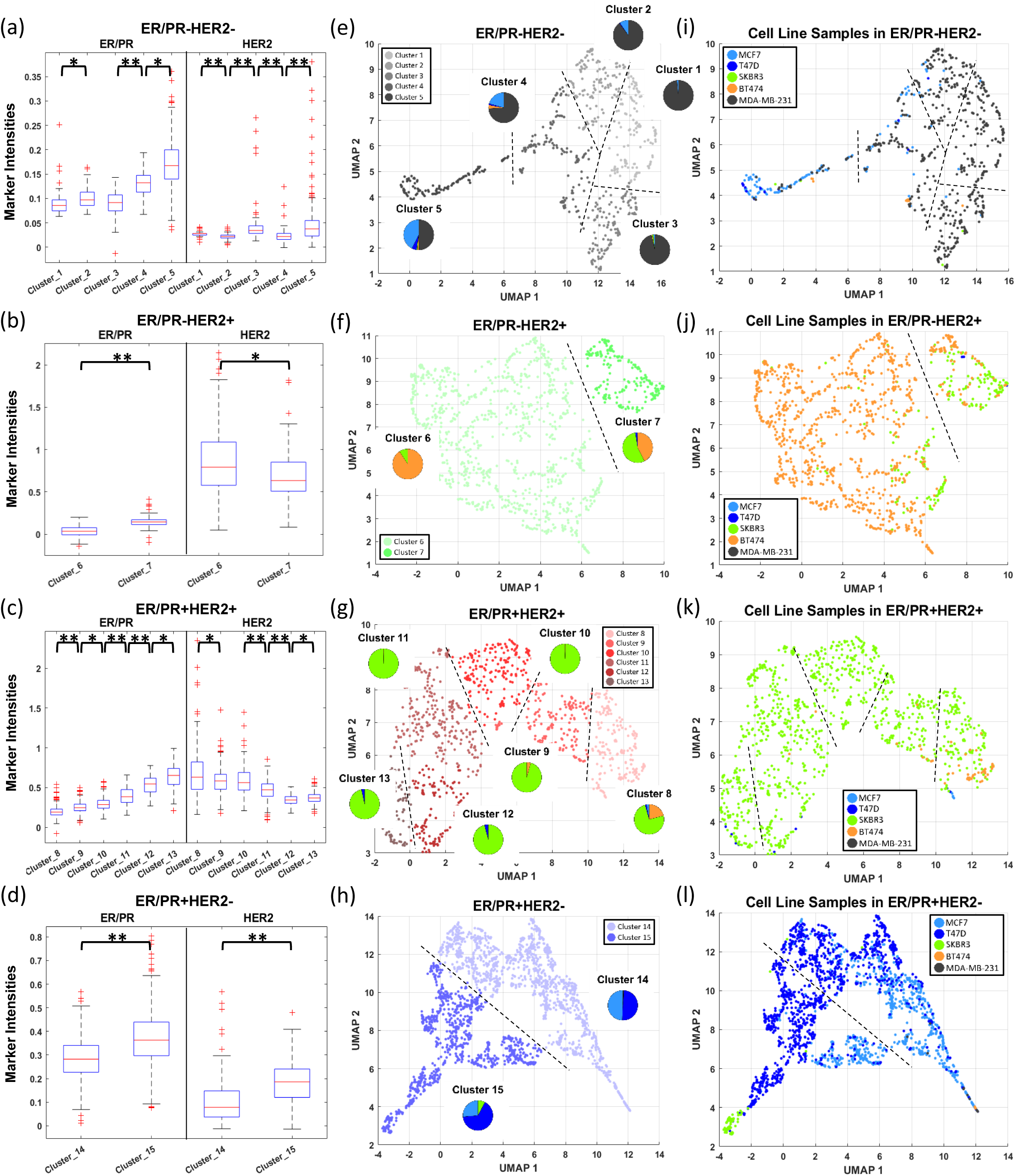
Characteristics of the identified subclusters from reconstructed images. **(a-d)** Differences of the mean intensity values of the subclusters. **(e-h)** UMAP visualization of reconstructed image features color-coded with the subclusters. The pie plots describe the proportion of the cell lines. (**i-l**) UMAP visualization of reconstructed image features color-coded with the cell lines. *p=10^- 10^∼10^-2^ and **p<10^-10^ .

**Supplementary Figure 5.**
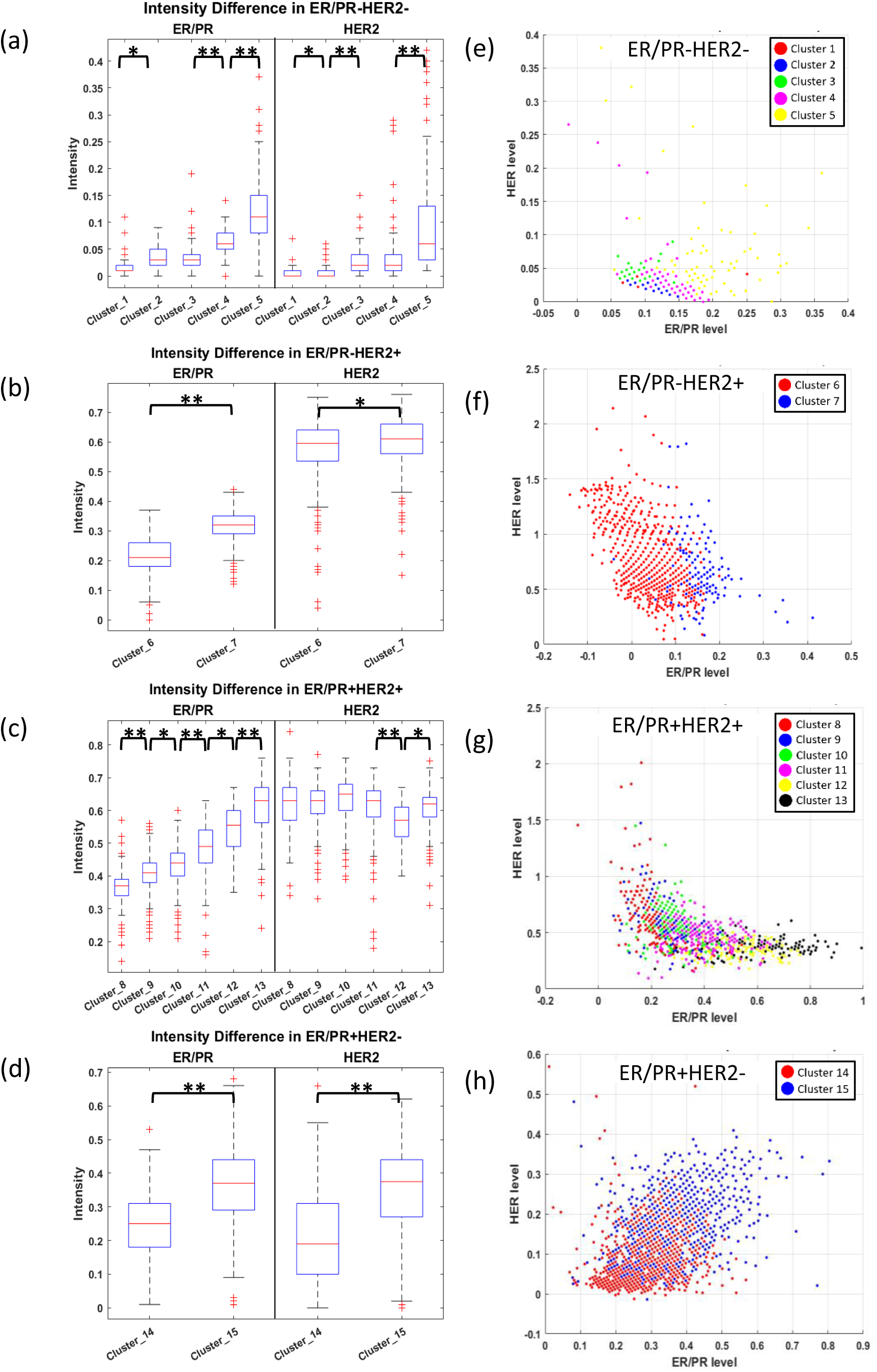
Characteristics of the identified subclusters of breast cancer cells from reconstructed images. **(a-d)** Differences of the mean raw intensity values of the subclusters in each breast cancer cell type. **(e-h)** Distribution of marker intensities in each breast cancer cells color-coded with the subclusters. *p=10^-10^∼10^-2^ and **p<10^-10^ .

**Supplementary Figure 6.**
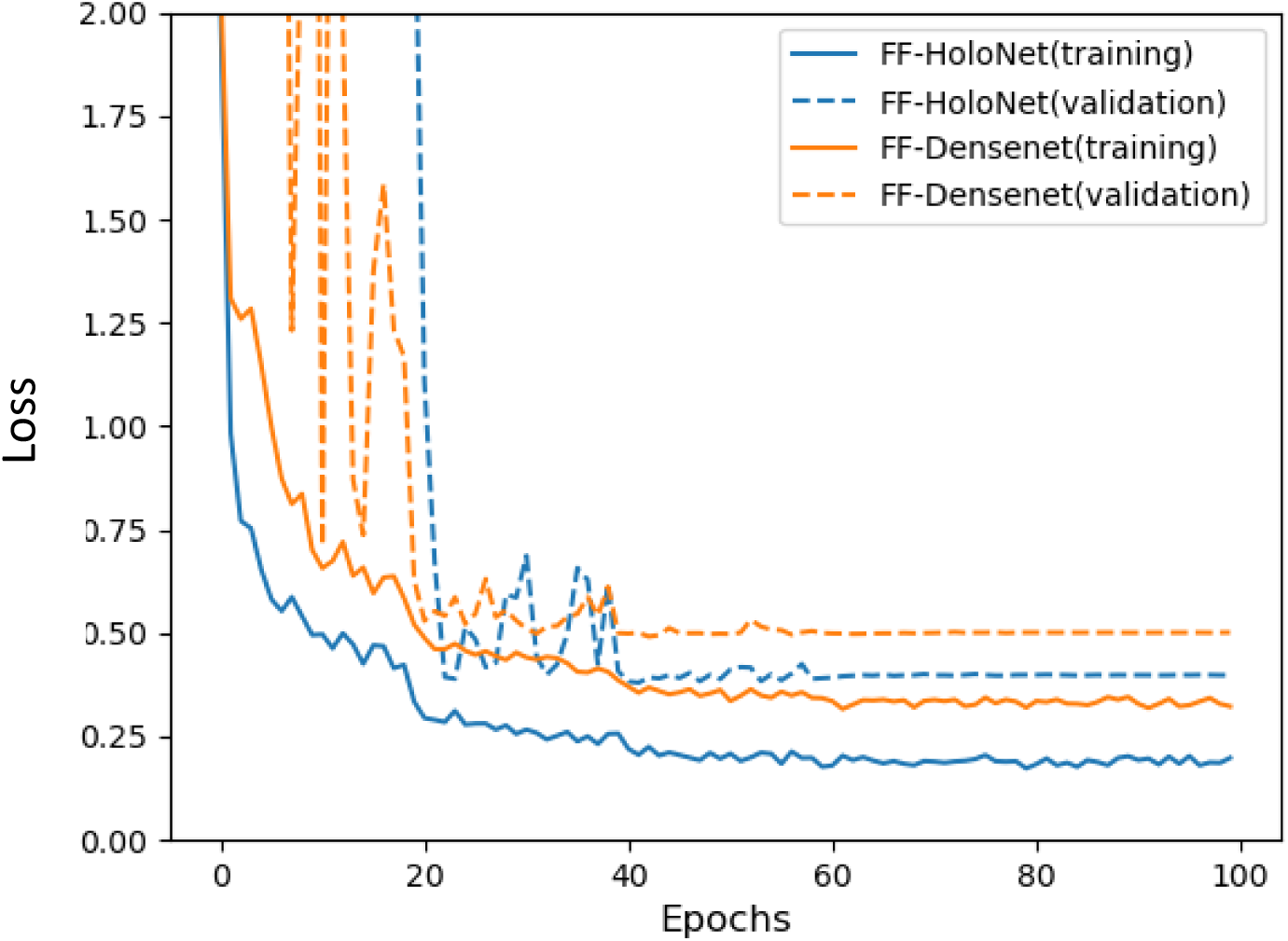
Training and validation losses of feature-fusion (FF) HoloNet and FF DenseNet model.

**Supplementary Figure 7.**
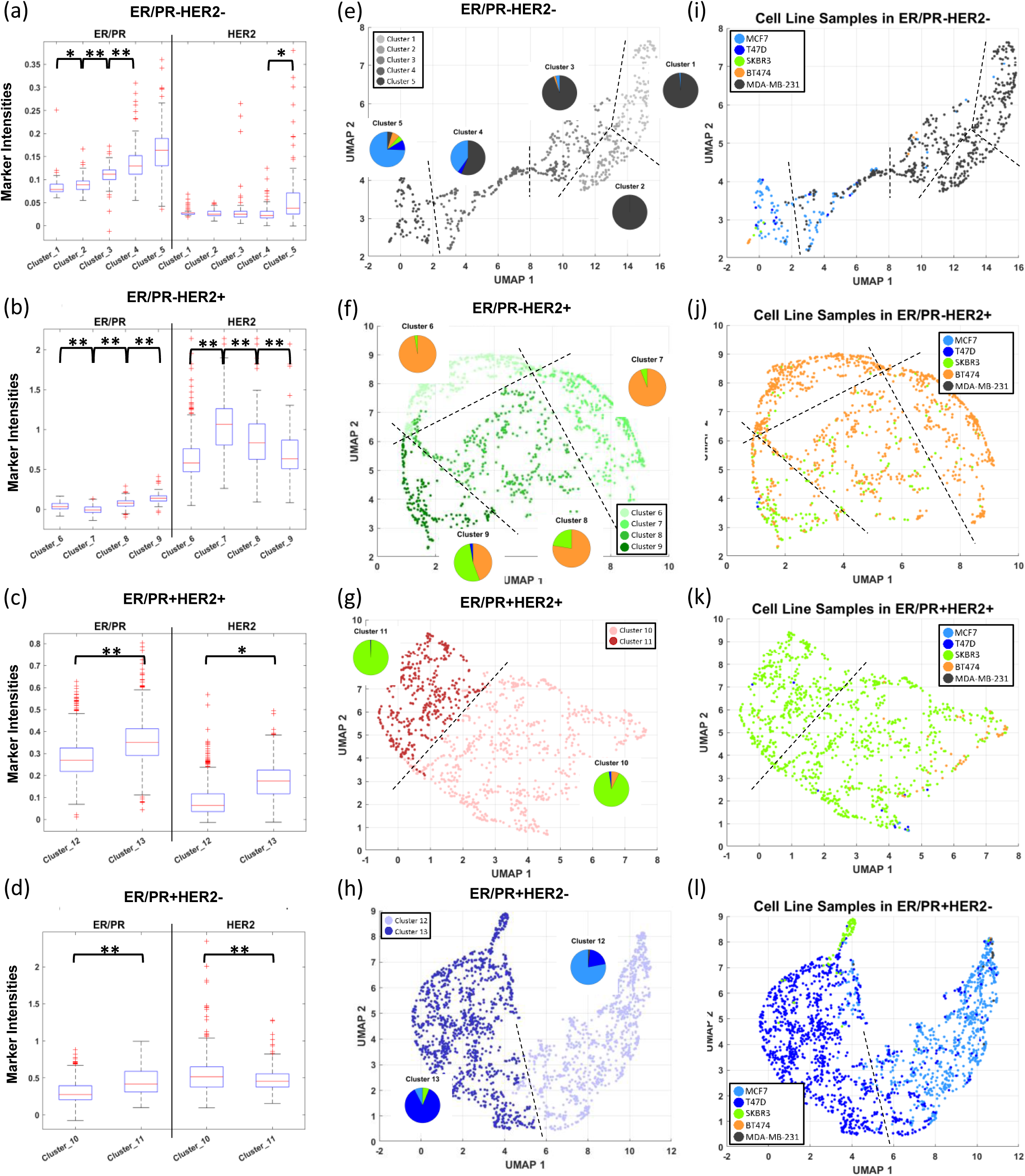
Characteristics of the identified subclusters extracted from the feature-fusion DenseNet model. **(a-d)** Differences of the mean intensity values of the subclusters. **(e-h)** UMAP visualization of hologram features color-coded with the subclusters. The pie plots describe the proportion of the cell lines. (**i-l**) UMAP visualization of hologram image features color-coded with the cell lines. *p=10^-10^∼10^-2^ and **p<10^-10^ .

**Supplementary Table 1.** The comparison of the classification performance among different models. The F1-score of each breast cancer cell type are also presented.

| Type | Accuracy | F1-Score |  |  |  |  |
| --- | --- | --- | --- | --- | --- | --- |
|  |  | ER/PR-HER2- | ER/PR-HER2+ | ER/PR+HER2+ | ER/PR+HER2- | Total |
| Previous Model | 0.902 | - | - | - | - | - |
| CNN | 0.9397±0.0031 | 0.941±0.004 | 0.9508±0.0049 | 0.9346±0.0041 | 0.9308±0.0037 | 0.9393±0.0042 |
| ResNet | 0.9362±0.0029 | 0.9407±0.0043 | 0.9498±0.0017 | 0.9253±0.0037 | 0.927±0.0044 | 0.9357±0.0035 |
| DenseNet | 0.9537±0.0024 | 0.9545±0.0033 | 0.9654±0.0011 | **0.9485±0.0025** | 0.945±0.0044 | 0.9533±0.0028 |
| HoloNet Model (24X24) | 0.9522±0.0023 | 0.9533±0.0061 | 0.9465±0.0061 | 0.9248±0.0054 | **0.9695±0.0014** | 0.9522±0.0023 |
| HoloNet Model | **0.9549±0.0025** | **0.956±0.0033** | **0.9707±0.0026** | 0.9467±0.0052 | 0.9437±0.0073 | **0.9553±0.0046** |

## Notes

### Competing Interest Statement

The authors have declared no competing interest.

### Summary of Updates

Figure 2 revised; new Figure 3 added

